# Alternative splicing QTLs in European and African populations using Altrans, a novel method for splice junction quantification

**DOI:** 10.1101/014126

**Authors:** Halit Ongen, Emmanouil T. Dermitzakis

## Abstract

With the advent of RNA-sequencing technology we now have the power to detect different types of alternative splicing and how DNA variation affects splicing. However, given the short read lengths used in most population based RNA-sequencing experiments, quantifying transcripts accurately remains a challenge. Here we present a novel method, Altrans, for discovery of alternative splicing quantitative trait loci (asQTLs). To assess the performance of Altrans we compared it to Cufflinks, a well-established transcript quantification method. Simulations show that in the presence of transcripts absent from the annotation, Altrans performs better in quantifications than Cufflinks. We have applied Altrans and Cufflinks to the Geuvadis dataset, which comprises samples from European and African populations, and discovered (FDR = 1%) 1806 and 243 asQTLs with Altrans, and 1596 and 288 asQTLs with Cufflinks for Europeans and Africans, respectively. Although Cufflinks results replicated better across the two populations, this likely due to the increased sensitivity of Altrans in detecting harder to detect associations. We show that, by discovering a set of asQTLs in a smaller subset of European samples and replicating these in the remaining larger subset of Europeans, both methods achieve similar replication levels (94% and 98% replication in Altrans and Cufflinks, respectively). We find that method specific asQTLs are largely due to different types of alternative splicing events detected by each method. We overlapped the asQTLs with biochemically active regions of the genome and observed significant enrichments for many functional marks and variants in splicing regions, highlighting the biological relevance of the asQTLs identified. All together, we present a novel approach for discovering asQTLs that is a more direct assessment of splicing compared to other methods and is complementary to other transcript quantification methods.

## Introduction

In eukaryotes alternative splicing is involved in development, differentiation (Graveley et al. 2011) and disease (Yoshida et al. 2011) in a tissue specific manner. Splicing events can be categorized under skipped exon, retained intron, alternative 3’ or 5’ splice sites, mutually exclusive exons, alternative first or last exons, or tandem UTR categories. Before the invention of microarray technology the proportion of multi-exonic genes undergoing alternative splicing was estimated at approximately 50% (Lander et al. 2001). However as the technology improved these estimates increased to 74% with microarrays (Johnson et al. 2003) and to almost 100% with RNA-sequencing (Wang et al. 2008). Although RNA-sequencing has been a very powerful tool in discovering unique transcription in tissues and diseases (Wang et al. 2012), and also in elucidating the regulation of transcription (Montgomery et al. 2010; Pickrell et al. 2010; Lappalainen et al. 2013), accurately quantifying transcripts remains a challenge due to the short read length used in most population based studies. Currently there are multiple transcript quantification methods available including *de novo* quantification methods like Cufflinks (Trapnell et al. 2010) and Scripture (Guttman et al. 2010), and annotation-based methods like MISO (Katz et al. 2010) and Flux Capacitor (Montgomery et al. 2010). However both approaches have inherent flaws since *de novo* methods make the assumption that the most parsimonious solution best describes the underlying transcriptome and annotation based methods assume complete knowledge of the transcriptome, both of which are unlikely to be true.

In this study we present a novel method for relative quantification of splicing events from RNA-sequencing data called Altrans. Our approach is an annotation-based method, which makes the least number of assumptions from the annotation. To this end we chose to simplify the problem and quantify relative frequencies of observed exon pairings in RNA-sequencing data for all categories of splicing events. This approach only assumes correct knowledge of the exons in the transcriptome, and is agnostic to the isoform structures defined in an annotation, which would, in theory, make it more accurate and sensitive in the presence of unknown isoforms. We tested the performance of Altrans versus a well-established transcript quantification method, Cufflinks (Trapnell et al. 2010) and benchmarked our method in two ways. Firstly we conducted a simulation study and assessed the concordance of the measured quantifications by each method with the simulated quantifications. Secondly, we assessed the relative power of discovering alternative splicing quantitative trait loci (asQTLs) for each method. For the asQTL analyses we chose the Geuvadis dataset, since it was, at the time of analyses, the largest publically available population-based RNA-sequencing study. The Geuvadis dataset comprises 462 individuals in the 1000 Genomes project (Genomes Project et al. 2012) from 5 populations: the CEPH (CEU), Finns (FIN), British (GBR), Toscani (TSI) and Yoruba (YRI), and contains data for whole genome DNA sequencing and deep mRNA sequencing in the lymphoblastoid cell line (LCL) (Lappalainen et al. 2013), and is thus an ideal dataset for our purposes.

## Results

### Simulation results

The general overview of the Altrans algorithm is provided in Figure 1. We first aimed to compare the results between Altrans and Cufflinks using simulations. We compared 4 scenarios, one where the given annotation perfectly described the transcripts in the simulations and three others with 25%, 50%, and 75% novel transcripts absent from the annotation. Subsequently we quantified the 4 simulation results with both algorithms using the known annotation in all cases. This was done to assess how methods performed in cases of complete versus incomplete transcriptome knowledge. The transcript quantifications generated by Cufflinks were transformed into link quantifications to make them comparable to those generated by Altrans.

**Figure 1.**
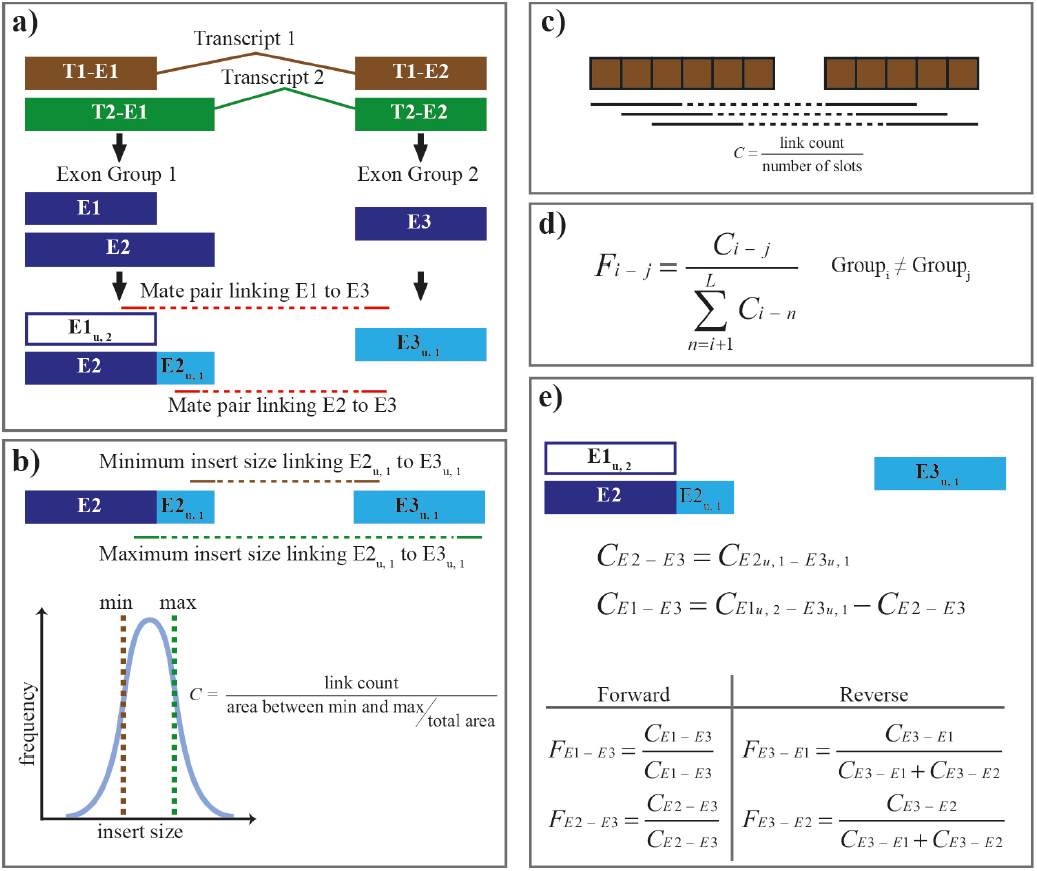
Schematic of the Altrans algorithm. (a) Overlapping exons are grouped into exon groups where identical exons belonging to multiple transcripts are treated as one unique entity. Two transcripts, shown as connected brown and green boxes, result in two exon groups and three exons shown as blue boxes. Next unique regions of each exon, depicted as light blue boxes and a subscript u followed by the level of the exon, are identified. Since E2 has a region which is not shared by any other exon it is assigned a “level” of 1, and the reads aligning to E2u,1 can be unambiguously assigned to E2. E1 does not have a unique portion, therefore the level 1 exon, E2, is removed from the exon group and the whole of E1 becomes a pseudo-unique portion, shown as an empty blue box, with a level of 2. These unique regions are used when assigning mate pairs to links as shown with the red lines where the solid portions of the line are the sequenced mates and the dashed part represents the inferred insert. (b) The default method for calculating link coverage. Link coverage is necessary to normalize the observed counts for the length of the unique portions being linked and the insert size. The theoretical minimum and maximum insert sizes linking the two unique portions, represented as brown and green lines respectively, are calculated and given the empirically determined insert size distribution, the area under the curve between the minimum and maximum insert sizes is estimated. The link coverage equals the number of mate pairs linking the two unique portions over the ratio of this area to the area of the whole insert size distribution. (c) The slots method for determining link coverage. Here given a read length and insert size of 3 and two exons that are 6 and 5 bases long, there are three slots, theoretical number of positions where a mate pair (given, in this case, 3+3+3 = 9 bp long fragment size) exists that links these exons on the mRNA, shown as black lines. The link coverage is the number of mate pairs linking the exons over the number of slots. (d) The equation to calculate F value for a link. (e) A worked example of calculation of the F values. First the coverage of E2 to E3 link (CE2-E3) is determined from level 1 unique regions (CE2u,1-E3u,1) which is then subtracted from the coverage attained from the pseudo-unique E1 to E3 link (CE1u,2-E3u,1) in order to calculate the true E1-E3 coverage (CE1-E3). In the forward direction E1 and E2 become primary exons and in the reverse direction E3 is the primary exon and the corresponding F values are calculated as shown.

The results of the simulation analysis are shown in Figure 2. We observe that Cufflinks performs better than Altrans when the annotation is perfect, but as the percentage of novel transcripts in the simulations increases Altrans performs better as it suffers less from the imperfect annotation used in the quantification. In order to produce a null random distribution for each method we took the link quantifications for each gene and permutated these for 100 times within the links of this gene. We then measured the correlation of these random assignments with the simulated ones and find that Cufflinks falls to the levels of random assignment of link quantifications as the novel transcripts increase in the simulations. We estimated the proportion of novel transcripts by using split read mappings from a well-studied LCL transcriptome RNA-sequencing experiment (Lappalainen et al. 2013) and a less well-studied pancreatic beta cell transcriptome RNA-sequencing experiment (Nica et al. 2013). We observe that in the LCLs on average 25.8% (sd = 3.5%) and in the beta cells 34.7% (sd = 9.3%) of the junctions are not found in the GENCODE version 12 annotation. Therefore we conclude that in RNA-sequencing experiments where the annotation does not fully reflect the underlying isoform variety Altrans is a sensitive method for quantifying exon junctions.

**Figure 2.**
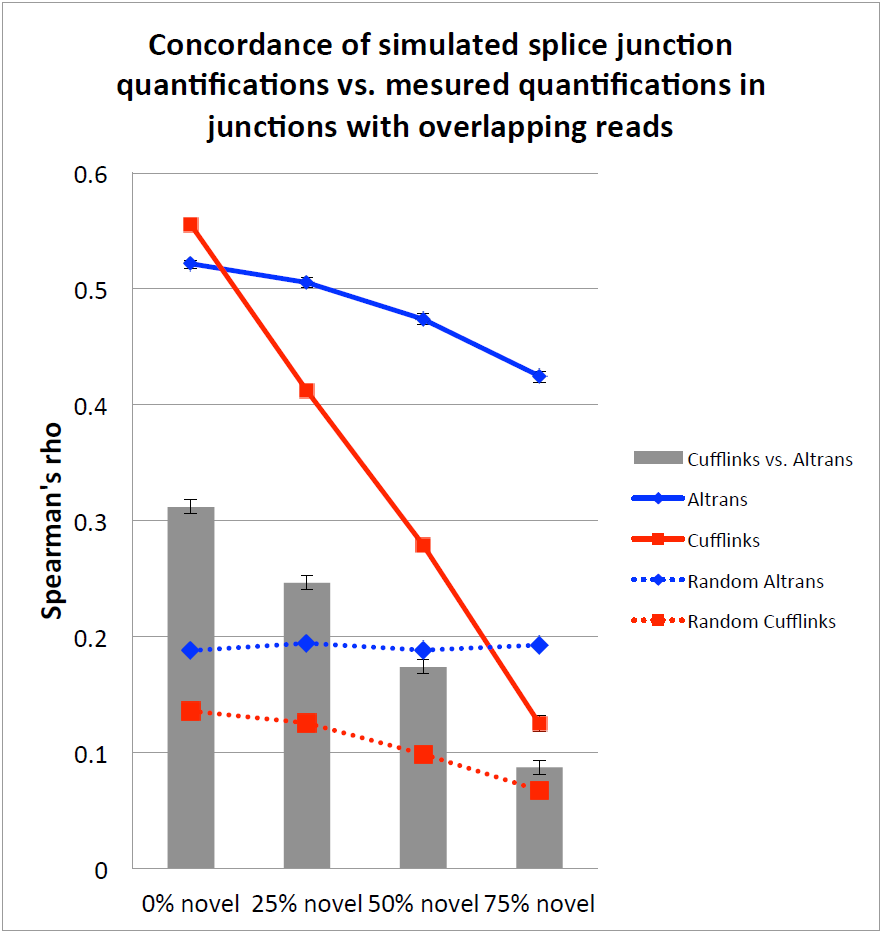
Simulation results. Using Flux Simulator we ran 4 simulations with varying levels of unannotated transcripts. Subsequently we ran quantifications with 2 methods with the known the GENCODE version 12 annotation. We compared the simulated vs. measured link quantifications using Spearman’s rank correlation. These comparisons are shown as colored solid lines. In order to produce a null random distribution for each method we took the link quantifications for each gene and permutated these for 100 times within the links of this gene and measured the correlation of these random assignments with the simulated ones. By using this sampling method stratified by genes we account for the variability of number of isoforms per gene. These correlations for random assignments are shown as dashed lines. We also compared the concordance of Altrans with Cufflinks shown as the grey bars. We observe that as the percentage of novel transcripts increase the performance of Cufflinks suffers, whereas this is not the case for Altrans, which results in best quantifications with increased levels of unannotated transcripts.

### Cis-aiternative splicing QTL discovery and replication between populations

The Geuvadis dataset comprises 373 European (EUR) and 89 African (YRI) samples and the cis-asQTL discovery was conducted separately in each population as described in the methods section. At an FDR threshold of 1% we find 1806 and 1596 asQTL genes in the European population with Altrans and Cufflinks, respectively. For the Africans these numbers are 243 and 288, respectively (Table 1). There is a significant overlap between the methods in the asQTL genes, with Altrans finding approximately 53% of the genes identified by Cufflinks in the Europeans and about 35% in the Africans (Table 1). The relative decrease of overlap between the methods in the African population is due to the decreased samples size, hence power, in this cohort compared to the Europeans. When we plot the significant asQTLs distances from the transcription start site (TSS) we observe that for both methods the asQTLs that are shared between the two populations and asQTLs with stronger effects tend to be closer to the TSS than population specific and weaker asQTLs (Figure 3a). As expected, given the sample sizes of each population, majority of the asQTLs in Europeans at this FDR threshold are unique to this populations (90% for Altrans and 85% for Cufflinks) whereas most of the African asQTLs are also found in the Europeans (78% for Altrans and 83% for Cufflinks) (Figure 3b). Using a more sensitive π_1_ approach (Storey and Tibshirani 2003) we estimate that 53% of the Altrans asQTLs in Europeans are replicated in Africans and 97% of the African asQTLs are replicated in Europeans. In the case of Cufflinks these estimates are 69% and 97%, respectively (Figure 3c).

**Figure 3.**
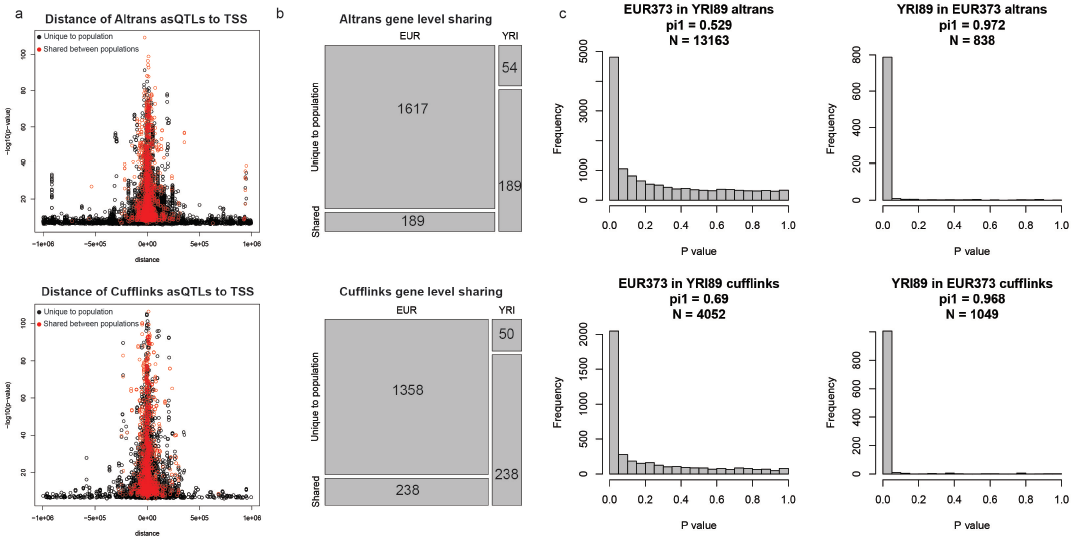
asQTL discovery. (a) The relative distance of asQTLs to the TSS versus the p-value. (b) Mosaic plots of gene level sharing of asQTLs for each method at FDR = 1%. (c) The p-value distributions of a variant-link pair tested in the other population for each method. From these p-value distributions the π_1_ statistic is calculated which estimates the proportion of true positives.

**Table 1.** Number of genes tested and asQTLs discovered at FDR = 1% in each population and by each method. The overlap column lists the common genes between the methods and the p-value refers to this overlap arising by chance.

| Population | Number of genes Altrans | asQTLs Altrans | Number of genes Cufflinks | asQTLs Cufflinks | Overlap | Overlap P-value |
| --- | --- | --- | --- | --- | --- | --- |
| EUR | 7443 | 1806 | 7148 | 1596 | 814 | 8.9e-157 |
| YRI | 7720 | 243 | 7391 | 288 | 100 | 3.97e-83 |

We have taken the Spearman’s rho as a proxy to the effect size of an asQTL, and compared the absolute value distribution of the rhos of significant asQTLs identified by each method both populations (**Supplementary Figure 1**). Cufflinks asQTLs’ have significantly higher effect sizes than Altrans asQTLs (Mann-Whitney *P* < 2e-16 in Europeans and *P* < 5.4e-10 in Africans) in both populations, indicating that Altrans is identifying associations with smaller effect sizes compared to Cufflinks and together with changes in sample size, this contributes to relative lack of replication of Altrans asQTLs across populations. In order to test the replication of asQTLs by each method independent of sample size and different populations, we have selected 91 European individuals belonging to the CEU population and replicated the findings of this cohort in the larger 282 remaining European samples. When we calculate the π_1_ statistic in this analysis we observe the both methods attain similar levels of replication (π_1_= 94% for Altrans and 98% for Cufflinks) (**Supplementary Figure 2**). In the absence of a known “true” set of asQTLs we have taken the number of asQTLs discovered as a proxy to the sensitivity of a method, since there is no prior to suggest that genotypes would randomly correlate with splicing. Moreover, we used the π_1_ values attained from the replication within the European samples and the total number of genes tested by each method to correct the number of significant asQTLs identified by each method (Methods). We observe that, when these corrections are applied, Altrans finds 1630 asQTLs in Europeans and 219 asQTLs in Africans whereas Cufflinks identifies 1564 and 282 asQTLs in Europeans and Africans, respectively, which are highly likely to be true positives. This similar levels of sensitivity achieved by each method indicates the validity of both approaches.

### Replication of discoveries by one method in the other method

We wanted to assess how discoveries of one method compared to the other. For each significant variant-link pair in one population by one method, we calculated the p-value of the same variant-link pair in the same population based on quantifications by the other method. From these p-value distributions we calculated the π_1_ statistic, which indicates the proportion of true positives (**Supplementary Figure 3**). We estimate that 49% of Altrans asQTLs in Europeans and 52% Altrans asQTLs in Africans are replicated by Cufflinks quantifications in the corresponding population. Whereas replication in the other direction, Cufflinks asQTLs in Altrans, is higher and is 78% and 90% for Europeans and Africans, respectively. This suggests that asQTL discovered by Cufflinks are common between the methods more so than the asQTLs found by Altrans, confirming the complementary nature of the two approaches.

Moreover we compared the types of splicing events that are found to be significant by both methods (**Supplementary Figure 4**) and observed that there are significant differences between the two methods. The majority (66%) of the signal that Altrans captures is due to exon skipping events followed by alternative 5’ and 3’ UTRs (15% and 11%, respectively). In comparison Cufflinks has a more uniform distribution of significant event types with the most common being alternative 5’ UTR (23%), followed by exon skipping (15%) and alternative first exons (14%). This difference in types of significant splicing events each method finds, highlights their relative merits in identifying different types of splicing events and is the main reason for method specific significant results.

Although the methodology in identifying splicing QTLs in the original Geuvadis analysis differs significantly from the process described here, we also checked the asQTL gene level overlap between the published lists of splicing QTLs (Lappalainen et al. 2013) and the ones identified here (**Supplementary Figure 5**). We find that Altrans detects 283 out of the 620 asQTLs identified in the Europeans in the original study, and Cufflinks finds 333 overlapping asQTLs. The union of both methods used here identifies 395 genes as significant asQTLs out of the 620 in the original discovery. In the African population the overlap proportions are similar with Altrans finding 27 out of 83 asQTLs as also significant, whereas Cufflinks finds 34 common genes, and the union of Altrans and Cufflinks overlaps with 41 asQTLs in the original study. This is another confirmation of the complementary nature of asQTL discovery methods.

### Functional relevance of asQTLs

In the absence of a known and true set of asQTLs we can use the functional annotation of the human genome generated by the ENCODE project to assess whether the asQTLs discovered are likely to be biologically active. If the identified asQTLs are “real” then we would expect them to lie in biochemically functional regions of the genome more often than expected by chance. We have tested this by overlapping asQTLs with functional annotations provided by the Ensemble regulatory build (Flicek et al. 2013) and comparing this overlap to that of random set non-asQTL variants, which were matched to the asQTLs based on relative distance from TSS and allele frequency (Methods). We find significant enrichments for many transcription factor peaks (median 2.7 × median *P* = 9.49e-8 for Altrans and median 3.3 × median *P* = 1.51e-7 for Cufflinks), DNase1 hypersensitive sites (2.4 × *P* = 2.57e-32 for Altrans and 2.0 × *P* = 2.31e-14 for Cufflinks), chromatin marks for active promoters (median 2.6 × median *P* = 2.05e-37 for Altrans and median 2.7 × median *P* = 3.64e-37 for Cufflinks) as well as strong enhancer marks (median 2.3 × median *P* = 9.36e-87 for Altrans and median 2.3 × median *P* = 2.32e-16 for Cufflinks) in asQTLs identified by both methods (Figure 4). We also observe a significant depletion in repressor marks (3.4 × *P* = 1.56e-48 for Altrans and 4.4 × *P* = 1.60e-42 for Cufflinks). All together these results confirm the functional relevance of asQTLs and indicate that we are capturing true biological signal. Furthermore we also observe strong significant enrichments for variants that are in splice acceptor (17.1 × *P* = 7.95e-9 for Altrans and 33.3 × *P* = 7.95e-9 for Cufflinks) and donor sites (3.3 × *P* = 0.05 for Altrans and 6 × *P* = 0.03 for Cufflinks) as well as variants in splice regions (3.4 × *P* = 3.9e-9 for Altrans and 7.6 × *P* = 1.99e-26 for Cufflinks), which also indicates that we are capturing variants involved in splicing machinery.

**Figure 4.**
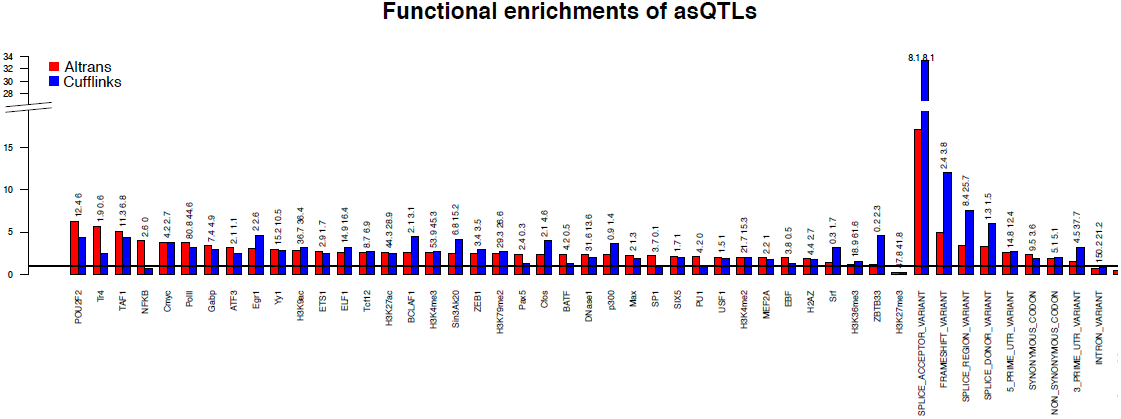
Figure 4. Functional enrichments of asQTLs discovered by Altrans (red) and Cufflinks (blue). All variants identified in separate populations are merged. The null (frequency and distance matched) is represented as the black horizontal line. The numbers above each bar are the-loglO p-values of the enrichment, Altrans enrichment p-value followed by Cufflinks p-value.

## Discussion

Here we present a novel method, Altrans, for relative quantification of splicing events (Figure 1) to be used in population genetics studies in discovery of asQTLs. Since the phenotype is splicing ratios of exon links calculated from mapping of RNA-sequencing reads without modeling of transcript structure, it is a more direct estimation of splicing. We have assessed the performance of the Altrans algorithm versus the Cufflinks method both on simulated and biological data. The simulation analysis indicates that when the annotation perfectly describes the underlying isoform variety Cufflinks performs better than Altrans. This is due to the fact that Cufflinks quantifies transcript rather than exon links, i.e. it uses the total length of the transcript in quantifications, whereas Altrans only uses the observed reads that are linking a pair of exons. When we convert the transcript quantifications of Cufflinks into link quantifications, this means that all the links in a transcript will “borrow” information from other links of the transcript; whereas in Altrans all the links will be independently measured from the observed reads overlapping the link. Moreover, when the perfect annotation is available using transcript quantifications, as in the case of Cufflinks, is a more accurate approach. However the simulations also show that when there are novel transcripts, i.e. isoforms that are not explained by the annotation, the accuracy of transcript quantifications decreases for Cufflinks whereas Altrans quantifications do not suffer as much as the transcript quantifications. We estimate that in less well-studied transcriptomes like the human pancreatic beta cell transcriptome (Nica et al. 2013) the proportion of the links between exons that are novel would be high enough that using the known annotation may result in unreliable estimates.

It is important to assess the performance of a method using biological data, and we applied Altrans and Cufflinks to the Geuvadis dataset (Lappalainen et al. 2013) with the specific aim of identifying asQTLs. We find 1806 and 1596 asQTL genes in the European population and 243 and 288 asQTLs in the Africans with Altrans and Cufflinks, respectively. There is a decrease in the replicated variant-link pairs between populations in Altrans when compared to Cufflinks. However this is due to Altrans detecting harder to detect associations, reflected in the effect sizes of the asQTLs identified by each method (**Supplementary Figure 1**). Using two subsets from the European samples we show that Altrans and cufflinks achieve similar levels of replication. The method specific asQTLs are explained by the different types of alternative splicing events each method captures (**Supplementary Figure 4**). Altrans is more powerful in capturing exon skipping events, whereas Cufflinks appears to be as powerful in capturing events in the ends of transcripts. This is an expected result given how each method works. Since Altrans is examining reads that link multiple exons it will perform relatively poorly when a read pair has to extend over constitutive parts of exon groups if constitutive parts are larger than the insert size of the experiment, since there will be very few reads joining these types of exons. On the other had since Cufflinks uses all reads over a transcript it will not fail to quantify these types of event accurately and this is reflected in the types of events each algorithm identifies. Furthermore, when we compare replication of results of one method by the other method we observe that Cufflinks results replicate better in Altrans than in the other direction (**Supplementary Figure 3**). These results taken together indicates that Altrans has increased sensitivity over Cufflinks in detecting certain types of splicing event.

The relevance of these asQTLs identified by both methods is confirmed by their significant overlap with functional annotations. This result, in the absence of a comprehensive list of asQTLs, shows that asQTLs that we are capturing reside in biochemically active regions of the genome, which reaffirms that we are capturing real biological signal. However there are differences in the asQTLs detected by different methods (**Supplementary Figure 5**). This is due to the fact that in RNA-sequencing with short read lengths, all methods have to infer quantifications of transcript or splice junctions, and each method in doing so has its relative merits. Here we present a novel approach to this problem, called Altrans, and show that it is sensitive and performs comparably to other methods. The Altrans software is available at https://sourceforge.net/projects/altrans/. We believe it will prove useful in the search for alternative splicing QTLs in population genetics studies.

## Methods

### Altrans method for relative quantification of splicing events

Altrans is a method for the relative quantification of splicing events. It is written in C++ and requires a BAM alignment file (Li et al. 2009) from an RNA-seq experiment and an annotation file in GTF format containing exon locations. The BAM file is read using the BamTools API (Barnett et al. 2011). Altrans utilizes paired end reads, where one mate maps to one exon and the other mate to a different exon, and/or split reads spanning exon-exon junctions to count “links” between two exons. Only the primary alignments of reads aligning to multiple locations are considered. The first exon in a link is referred to as the “primary exon". The algorithm is as follows:

1. Group overlapping exons from annotation into exon groups. Since we are quantifying splicing events and not individual transcripts, transcript level information is ignored and exons with identical coordinates belonging to multiple transcripts are treated as one unique exon (Figure 1a).
2. In order to assign reads to overlapping exons, identify unique portion(s) of each exon in an exon group. Exons with immediate unique portions, where there is no other overlapping exon and where a read can be unambiguously assigned to the exon, are called “level 1 exons". For exons with no unique positions, remove the level 1 exons from the exon group to determine pseudo-unique positions for the remaining exons, where again there is no other overlapping exon after the removal of the level 1 exons, and increment the level of these exons. In the rare cases where an exon shares its start position with one exon and its end position with another, causing it to have no unique portion, then this exon is removed from the analysis in order to be able to assign unique portions to the remaining exons in the same exon group. Iterate through this process of removing exons that have unique or pseudo-unique regions until all exons in a group have unique or pseudo-unique portions (Figure 1a).
3. Use these unique and pseudo-unique portions to assign mate pairs or split reads to links (Figure 1a). Links assigned to pseudo-unique portions are putative assignments and “deconvolution” of these is handled in the next step.
4. Since exons with pseudo-unique portions share this region with other exon(s),reads aligning here may belong to multiple exons. In order to unambiguously quantify links between these exons we calculate “link coverage” for all pairs of exons in a given window size.The default method divides the link counts with the probability of observing such a link given the insert size distribution, which is empirically determined from pairs aligning to long exons (Figure 1b). The second method involves calculating the number of “slots” linking two exons given the empirically determined mostfrequent insert size (Figure 1c). The link coverage metric ensures that the link counts are normalized for the specific insert size distribution of the experiment, which has direct effect on the observed link counts. Hence linkcoverage allows us to quantify an exon link from the unique portions only, i.e. this value should be equivalent to the one we would calculate if we were able to measure the whole exon. The coverage between level 1 exons can be calculated directly using the unique portions, whereas links between higher-level exons are calculated by iteratively subtracting coverage of allthe other lower level links from the coverage of these links (Figure 1e).
5. Calculate the quantitative metric, F value, for one exon link as the coverage of the link over the sum of the coverages of all the links that the primary exon makes (Figure 1d & e). Using this fraction rather than link counts or coverage ensures that the metric is independent of global effects on gene expression.
6. Repeat step 5 in both 5’-to-3’ (forward) and 3’-to-5’ (reverse) directions to capture splice acceptor and donor effects respectively (Figure 1e).

The program also allows the user to calculate an F value from all the links that a primary exon makes regardless of the direction. Among with the F values, the raw link counts are also outputted which allow filtering of results eliminating low count links. These raw counts can also be normalized, and subsequently reread by the program to calculate the F values. Memory usage and speed heavily depend on the complexity of the annotation and the number of reads in the alignment file. For a sample alignment with 50 million reads and an annotation with 539748 unique exons Altrans ran for 20 minutes and consumed 784 MB of RAM on a single 2.2 GHz core under Linux.

### Conversion of transcript quantifications to link quantifications

We convert the transcript quantifications generated with Cufflinks to relative link quantifications. This is achieved by assigning the same quantification to all linked exons of a transcript based on the measured quantification of the said transcript. We then apply the same method of relative link quantification used in the Altrans algorithm, specifically steps 5 and6 in the previous section, to calculate the F value for all the links a primary exon makes.

### Simulation analysis

In order to benchmark the link quantifications generated by Altrans we conducted a simulation analysis using the Flux Simulator software (Griebel et al. 2012). We simulated an RNA-sequencing experiment with 50 million reads with the GENCODE version 12 annotation (Harrow et al. 2012) reflecting cases where we have a perfect annotation describing all the observed transcripts in the data. Additionally we introduced novel transcripts, made up of existing exons of a gene, into the annotation and simulated 3 cases with 50 million reads where the novel transcripts accounted for 25%, 50%, and 75% of all transcripts reflecting cases where the annotation is not perfect. Altrans and Cufflinks (Trapnell et al. 2010) were run on these 4 simulated datasets using the standard GENCODE version 12 annotation. In each simulation the “correct” quantification of a transcript is taken as the RNA molecule count that the Flux Capacitor used to simulate reads for a given transcript. We have converted these “correct” transcript quantifications and the measured transcript quantifications of Cufflinks to exon link quantifications, as described in the previous section, and correlated the simulated expected link quantifications with the measured link quantifications for the 2 programs in the 4 simulation scenarios; using links where there were overlapping reads or links that were quantified both in the simulation and the given program. We measured the concordance between the simulated and measured quantifications using Spearman’s correlation. The estimates of novel splicing in a dataset are done through counting the number of uniquely mapping split reads. We then take junctions that are represented by at least 8 split reads and check whether this junction is present in the annotation.

### Cis-alternative splicing QTL discovery by each method in the Geuvadis dataset

The RNA-seq reads were aligned to the human reference genome (GRCh37) using the GEM aligner (Marco-Sola et al. 2012) and alignments were filtered for properly paired and uniquely mapping reads (mapping quality greater than or equal to 150). The genotype data were filtered for variants with MAF < 5% and HWE *P* < 1e-6 for each population separately and were corrected for population stratification using the first three and two eigenvectors for Europeans and Africans, respectively (Lappalainen et al. 2013). The Altrans link counts were normalized using the first 15 principal components calculated from these link counts. We filtered the Altrans results for primary exons that made at least 10 links in 30% of the samples that originated from exon groups with at least 15 links in 80% of the samples and this filter was applied to each population separately. Cufflinks quantifications were run using the annotation with the –GTF option. In the case of Cufflinks the transcript quantifications were converted to link quantifications and we assessed links originating from the same genes where there were Altrans quantifications. The cis-window for asQTL discovery was 1 mb flanking the transcription start site of each gene. The observed nominal p-values were calculated by correlating the genotype and link quantifications using Spearman’s rank correlation. Subsequently we ran permutations for each link separately to assign empirical p-values to each link. The permutation scheme involved permuting each link quantification at least 100 times to a maximum of 100,000 times, correlating this permuted data with all the genotypes in the cis-window and counting the number of permutations where the best permuted p-value was more significant than the observed best nominal p-value. This was done until 100 more significant permutation p-values were achieved with a maximum of 100,000 permutations per link. The empirical p-value is calculated as the number of permutations with a more significant p-value than the best nominal p-value of a given link over the total number of permutation iterations. These empirical p-values are then corrected for multiple testing using the qvalue R package (Storey and Tibshirani 2003).

In order to calculate the relative power of each method in finding significant asQTLs we used the 91 CEU samples to discover asQTLs. We then replicated these asQTLs in other non-overlapping 282 European samples. From the resulting p-value distributions we calculated the π_1_ statistic. We multiplied the number of significant asQTLs discovered by each method with the π_1_ statistic of each method, to identify the number of genes that are highly likely to be true positive asQTLs in each method. We then multiplied the Cufflinks asQTL gene number with the ratio of genes tested in Altrans to the genes tested in Cufflinks to account for the fewer number of genes tested in Cufflinks. This resulted in an adjusted count of “true” asQTLs for each method.

### Classification of splicing events

The alternative splicing events were classified into 10 categories, namely alternative 3’ splice site, alternative 3’ UTR, alternative 5’ splice site, alternative 5’ UTR, alternative first exon, alternative last exon, mutually exclusive exon, skipped exon, tandem 3’ UTR, and tandem 5’ UTR. For more information on these events refer to (Wang et al. 2008). We then classify each primary exon into these classes based on all of the observed links of the primary exon. This means that a primary exon may be involved in multiple splicing events. From these classifications we then calculate the proportion of each splicing class in the pool of significant primary exons. This method of classification was chosen since each link quantification is dependent on the quantification of all the other links that a primary exon makes.

### Functional enrichment of asQTLs

To compare the asQTL variants to a null distribution of similar variants without splicing association, we sampled genetic variants in the same *cis*-window of 1 mb surrounding the TSS and matched them to alternative splicing variants with respect to relative distance to TSS (within 5 kb) and minor allele frequency (within 2%). The variant effect predictor (VEP) (Adzhubei et al. 2010) tool from Ensembl was modified to produce custom tag which were STOP_GAINED, SPLICE_DONOR, SPLICE_ACCEPTOR, and FRAME_SHIFT. This modified version of VEP was applied to the imputed genotypes using the GENCODE v12 (Harrow et al. 2012) annotation. To this we added information of overlap with chromatin states (Ernst and Kellis 2010) and the Ensembl regulatory build (Flicek et al. 2013) which constituted our functional annotation. The enrichment for a given category was calculated as the proportion between number regulatory associations in a given category and all regulatory variants over the same proportion in the null distribution of variants. The p-value for this enrichment is calculated with the Fisher exact test.

## Acknowledgements

This research is supported by grants from European Commission SYSCOL FP7 (UE7-SYSCOL-258236), European Research Council, Louis Jeantet Foundation, Swiss National Science Foundation, and the NIH-NIMH (GTEx). The computations were performed at the Vital-IT (https://www.vital-it.ch) Swiss Institute of Bioinformatics.

